# Viral shedding and transmission after natural infection and vaccination in an animal model of SARS-CoV-2 propagation

**DOI:** 10.1101/2021.05.11.443477

**Authors:** Caroline J. Zeiss, Jennifer L. Asher, Brent Vander Wyk, Heather G. Allore, Susan Compton

## Abstract

At present, global immunity to SARS-CoV-2 resides within a heterogeneous combination of susceptible, naturally infected and vaccinated individuals. The extent to which viral shedding and transmission occurs on re-exposure to SARS-CoV-2 after prior natural exposure or vaccination is an emerging area of understanding. We used Sialodacryoadenitis Virus (SDAV) in rats to model the extent to which immune protection afforded by prior natural infection via high risk (inoculation; direct contact) or low risk (fomite) exposure, or by vaccination, influenced viral shedding and transmission on re-exposure. On initial infection, we confirmed that amount, duration and consistency of viral shedding were correlated with exposure risk. Animals were reinfected after 3.7-5.5 months using the same exposure paradigm. Amount and duration of viral shedding were correlated with re-exposure type and serologic status. 59% of seropositive animals shed virus. Previously exposed seropositive reinfected animals were able to transmit virus to 25% of naive recipient rats after 24-hour exposure by direct contact. Rats vaccinated intranasally with a related virus (Parkers Rat Coronavirus) were able to transmit SDAV to only 4.7% of naive animals after a 7-day direct contact exposure, despite comparable viral shedding. Observed cycle threshold values associated with transmission in both groups ranged from 29-36 cycles, however observed shedding was not a prerequisite for transmission. Results indicate that low-level shedding in both naturally infected and vaccinated seropositive animals can propagate infection in susceptible individuals. Extrapolated to COVID-19, our results suggest that continued propagation of SARS-CoV-2 by seropositive previously infected or vaccinated individuals is possible.

## Introduction

Individual immunity to SARS-CoV-2 infection may be gained through natural infection or vaccination. As a route to achieving population-wide immune protection, the former route, otherwise known as infection-induced herd immunity, is widely regarded as an ineffective strategy [1]. This route achieves unpredictable immunity [2–5] and incurs substantial morbidity [6] and mortality [5,7]. Consequently, mass vaccination against SARS-CoV-2 is underway as the safest and most effective means of controlling the COVID-19 pandemic [8].

For either route, our understanding of the extent to which previously naturally exposed [9–11] or vaccinated [12] individuals can shed and transmit virus on re-exposure is just emerging. Immunologic heterogeneity following natural infection [10,11,13], variable efficacy of different vaccines [8], and as yet unclear duration of immunity [14–16] are critical determinants of the level of herd immunity needed to control COVID-19. Depending on these variables, herd immunity needed to control COVID-19 varies widely from 50-100% [3,8].

Using a rat model of population-wide COVID-19 transmission, we assessed viral shedding and disease transmission following re-infection in naturally exposed or vaccinated seropositive individuals. Sialodacryoadenitis Virus (SDAV) is a highly infectious rat betacoronavirus [17,18] that infects the upper respiratory tract [19], salivary and lacrimal glands [20,21] and lung [21–23]. Like SARS CoV-2 [24–28], SDAV infection can be transmitted by asymptomatically infected individuals [29] via airborne, direct contact, or fomite routes [17].

Beginning with a defined SDAV inoculum, we modeled heterogeneous viral exposure in a naturally infected population using a range of high (inoculation and direct contact) and low (fomite) risk exposures. Previously exposed animals were then re-exposed to SDAV to determine the role of naturally acquired immunity in subsequent viral shedding and transmission to naïve animals. Shedding and transmission from naturally infected re-exposed animals was compared to that in animals vaccinated with Parker’s Rat Coronavirus (RCV), a closely related coronavirus [30,31].

We used the two most widely used measures of SARS-CoV-2 surveillance, reverse-transcriptase polymerase chain reaction (RT-PCR) testing [32] and serology [33] to examine the relationships between exposure routes, seroconversion and viral shedding.

## Materials and Methods

### Virus amplification and quantification

SDAV was isolated at Yale in 1976. Stocks of SDAV were generated in L2p176 cells [34,35]. Briefly, confluent L2.p176 cells were pretreated for 1 hour with 75ug/ml of DEAE-D in 75% DMEM 25% L15 media. Cells were incubated with virus diluted in 75% DMEM 25% L15 with trypsin for 1 hour and cell/media was harvested at 3 days post-infection. Viral titers were determined by plaque assay [34]. Briefly, 6 well plates of L2p176 cells were pretreated with 75ug/ml DEAE-D in 75% DMEM/25% L15 media for 3 hours. Cells were rinsed with PBS and inoculated with 10-fold dilutions of virus in 75% DMEM/25% L15 with 75ug/ml DEAE-D and trypsin. One hour later, inocula was removed, cells were rinsed with PBS and were overlaid with 0.55% Seaplaque agarose/minimal media/trypsin. Three days post-inoculation, cells were fixed with formalin, agarose was removed and plaques were visualized with Giemsa.

### Animals and housing

Seven-week old female and male SAS outbred Sprague-Dawley rats (150-250g) were purchased from Charles River Laboratories (Wilmington, MA). Animals were housed (separated by sex) in Tecniplast (West Chester, PA) individually ventilated cages (GR900 for rats) that provide high microbial biocontainment. Sentinel animals were placed on each side of the rack and tested every 3 months for antibodies to rodent pathogens, including SDAV, with consistently seronegative results. Rooms were maintained at 72°C on an evenly split light cycle (7AM:7PM). Animals were housed on corncob bedding, had access to autoclaved pellets (2018S, Envigo, Somerset, NJ) and acidified water ad lib, and were acclimated for 5-7 days prior to infection. Animals were individually identified using ear tags. These were regularly inspected and replaced as needed. Viral inoculation and animal handling was performed in a Class II biosafety cabinet. All exposure groups were separated by sex. All animal work was conducted under an approved Yale Animal Use and Care Committee protocol. Yale University is accredited by the Association for Assessment and Accreditation of Laboratory Animal Care

### Anesthesia, viral inoculation and euthanasia

Rats were anesthetized briefly using the open drop method (isoflurane: propylene glycol 30% v/v). Intranasal inoculation of 2X10^e^4 plaque-forming units (pfu) SDAV or 1X10^e^3 pfu RCV in Dulbecco’s Modified Eagle Medium (DMEM) was performed in a total volume of 50μl per animal (25 μl per nostril). Animals were fully recovered within 2-3 minutes of inoculation. At the end of reinfection and transmission experiments, animals were euthanized using 70% carbon dioxide.

### Initial infection with SDAV (**Figure 1A**)

Four exposure groups were defined for initial SDAV infection:

a. <u>Inoculated rats (n=19, 47% female).</u> Animals were inoculated intranasally with 2X10^e^4 pfu SDAV in 50μl DMEM, followed by individual housing in a clean cage for 48 hours.
b. <u>Direct contact (n=31, 48% female):</u> Naïve animals (1-3 animals) were placed with one inoculated rat in a new clean cage 48 hours post inoculation. After 24 hours, exposed rats were separated from the inoculated rat placed in a new clean cage.
c. <u>Fomite contact animals (n=55; 53% female).</u> Naïve animals (2-3 animals) were placed in a dirty cage (containing contaminated bedding, furniture, food and water nipple) that had been inhabited by an inoculated rat for 48 days post inoculation. After 24 hours, exposed rats were relocated to a new clean cage in small groups of 2-3 animals (fomite cohabitation group; n=30; 53% female) or singly (fomite single group; n=25; 52% female).
d. <u>Mock (n=10):</u> A control group (n=10; 50% female) was inoculated with DMEM alone.

Oral swabs were taken on day 2, 3, 4, 7 and 10 post-exposure on all animals (thus inoculated rats began this process 96 hours before other exposure groups). Each animal was weighed on its day of viral exposure and on every day on which oral swabs were taken. Gloves and equipment were sterilized with 200ppm MB-10 between each animal. Health checks were performed daily for 10 days post-exposure. Animals were bled 6 weeks after their 10-day initial exposure period to assess seroconversion.

### Reinfection with SDAV (**Figure 1B**)

After initial exposure, animals assumed either seronegative (n=66) or seropositive status (n=40). Within these two serologically distinct groups, those rats that had originally received intranasal infection with SDAV (inoculated rats) were re-infected intranasally again with the same viral dose (n=18). A subset of seronegative rats (n=18) received intranasal inoculations to provide a source of infection for remaining animals. Remaining animals were randomly assigned direct contact, fomite contact-cohabitation, and fomite contact-singly housed groups for their second exposure. Exposure and testing paradigms were identical to those described for initial reinfection. Time between initial and second exposure ranged from 113-165 days. Animals were euthanized at 10 days post infection (dpi) and assessed for seroconversion.

### Assessing transmission of SDAV to naïve rats by previously SDAV infected rats (**Figure 2A**)

Our aim was to determine whether animals that had been previously exposed to a known dose of SDAV could transmit SDAV to naïve rats following re-exposure after 4-5 months. Inoculated rats that had received an initial dose of 2X10^e^4 pfu SDAV, and had subsequently shed virus and seroconverted, received a second similar intranasal inoculation 113-165 days later. These animals were placed in a clean cage with naïve rats 48 hours post inoculation (direct contact paradigm). After 24 hours, exposed naïve rats were separated from the inoculated rat and placed in a new clean cage. Body weights and oral swabs were taken on both groups of animals at 2, 3, 4, 7 and 10 dpi, followed by serologic testing of naïve animals.

### Modeling vaccination using RCV (**Figure 2 B, C**)

First, we performed a dose-finding study to assess the dose of Parker’s rat coronavirus (RCV) that would impart protection to subsequent challenge with SDAV (**Supp Data 1**). Three groups of male rats (n=3/group) were inoculated with 10^e^3, 10^e^4 or 10^e^5 pfu RCV in 50 μl of DMEM. Seroconversion was confirmed in all 2 weeks after inoculation. Two weeks later, all animals were challenged with intranasal 2X10^e^4 pfu SDAV (50 μl). Low viral shedding (Cq 30.5-30.7 cycles for one day only over a 10-day period) was noted in only two animals at the higher RCV inoculation groups (10^e^4 or 10^e^5 pfu RCV). Based on these data, a dose of 10^e^3 RCV pfu RCV in 50 μl of DMEM was selected for the vaccination study. Eight-week-old naïve male and female SD rats were inoculated intranasally either once (n=12; **Figure 2B**) or twice with a month interval between inoculations (n=12; **Figure 2C**). Seroconversion was confirmed one month after the first inoculation.

**Figure 1.**
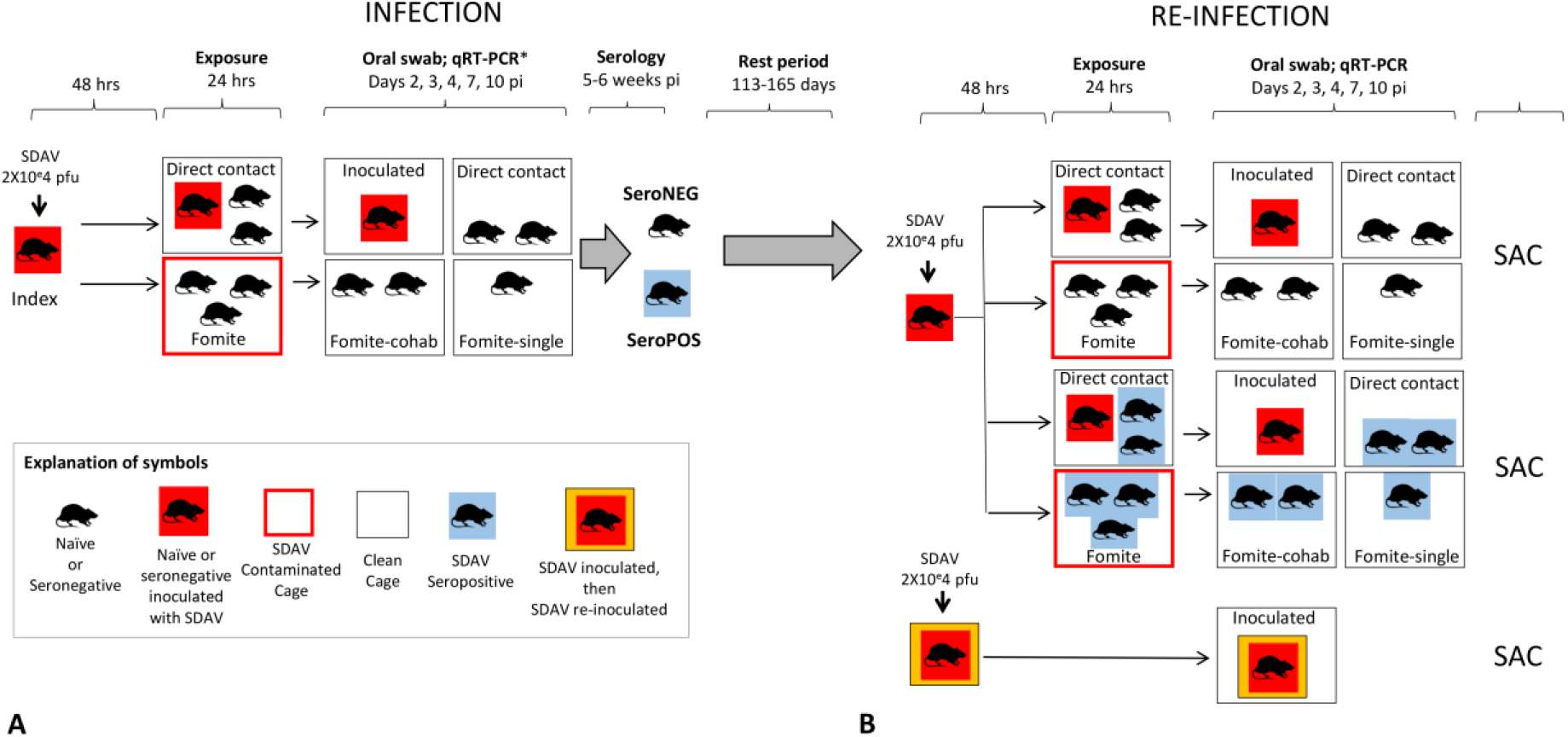
A. **Initial infection with SDAV:** Naïve animals were inoculated intranasally with 2X10^e^4 pfu SDAV. After 48 hours, one inoculated (index) animal was placed in a clean cage with naïve animals for 24 hours (direct contact)._Naïve animals were placed in a dirty cage that had been inhabited by an index rat (fomite) for 24 hours. After 24 hours, all rats were relocated to new clean cages to constitute four groups: index, or inoculated rats, direct exposure rats, and two fomite exposure groups – a fomite cohabitation group constituting 2-3 animals, and a fomite single group with only one animal. Oral swabs were taken on Day 2, 3, 4, 7 and 10 post-exposure on all animals, and serology performed 5-6 weeks later. A control group (not shown) was inoculated with DMEM alone, and similarly swabbed and bled. After initial exposure, animals assumed either seronegative or seropositive status. All groups were evenly split by sex. B. **Reinfection with SDAV:** Naïve seronegative rats received intranasal 2X10^e^4 pfu SDAV to provide a source of infection. Seropositive and seronegative animals were randomly assigned direct contact, fomite contact-cohabitation, and fomite contact-singly housed contact groups for their second exposure (Table 2 and Figure 1B). Rats that had originally received intranasal infection with SDAV (index rats) were re-infected intranasally again with the same viral dose. Time between initial and second exposure ranged from 113-165 days. Oral swabs were taken on Day 2, 3, 4, 7 and 10 post-exposure on all animals. Animals were sacrificed at 10 dpi and assessed for seroconversion. All groups were evenly split by sex.

**Figure 2.**
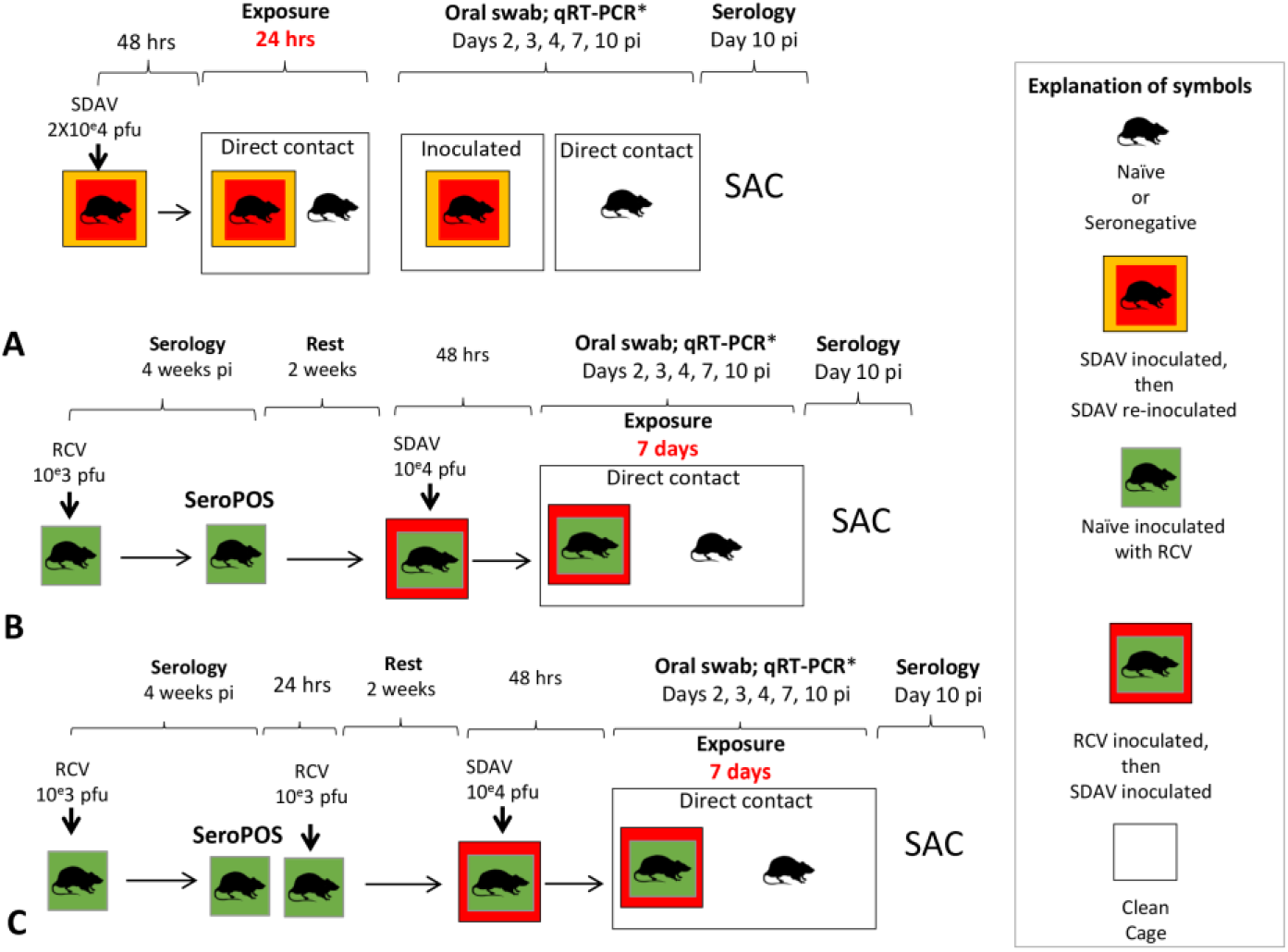
**A. Transmission of SDAV to naïve rats by previously SDAV infected rats**: Index rats that had received an initial dose of 2X10^e^4 pfu SDAV, and had subsequently shed virus and seroconverted, received a second similar intranasal inoculation 112-140 days later (n=12). After 48 hours, these animals were placed in a clean cage with naïve rats (n=12; direct contact paradigm). After 24 hours, exposed naïve rats were separated from the inoculated rat and placed in a new clean cage. Body weights and oral swabs were taken on both groups of animals at 2, 3, 4, 7 and 10 dpi. Animals were sacrificed at 10 dpi and assessed for seroconversion. All groups were evenly split by sex. **B. Transmission of SDAV to naïve rats by RCV (1 dose) vaccinated rats:** Eight-week-old naïve rats (n=12) were inoculated intranasally with 1X10^e^3 pfu RCV and assessed for seroconversion 4 weeks later. Two weeks after confirmed serologic status, animals were challenged with 2X10^e^4 pfu SDAV. After 48 hours, these animals were co-housed in a clean cage with a naïve rat (n=12) for 7 days. Body weights and oral swabs were taken on both groups of animals at 2, 3, 4, 7 and 10 dpi. Animals were sacrificed at 10 dpi and assessed for seroconversion. All groups were evenly split by sex. **C. Transmission of SDAV to naïve rats by RCV (2 doses) vaccinated rats:** The identical procedure was followed in this group with the addition of an additional RCV inoculation at the same dose 4 weeks after the initial inoculation 1X10^e^3 pfu RCV. Vaccinated rats (n=12); naïve recipients (n=9).

### Assessing transmission of SDAV to naïve rats by RCV vaccinated rats (**Figure 2 B, C)**

Next, we tested whether animals that had been previously vaccinated with RCV could transmit SDAV to naïve rats at all. Rats vaccinated with one or two doses of RCV were challenged by intranasal inoculation with 2X10^e^4 pfu SDAV in 50 μl of DMEM 6 weeks after their last RCV exposure. After 48 hours, one SDAV-inoculated rat was co-housed with one naïve rat for 7 days. Body weights and oral swabs were taken on both groups of animals at 2, 3, 4, 7 and 10 dpi, followed by serologic testing of naïve animals at 10 dpi. Animals were evenly allocated by sex.

### Assessing oral shedding by semi-quantitative RT-PCR

Rats were swabbed orally using sterile flocked swabs (Hydraflock, Puritan Medical Products, Guilford, ME) to confirm viral shedding by quantitative RT-PCR. 600 ul of RLT buffer (Qiagen) was added to each swab and the sample was vortexed. 350 ul of 70% ethanol was added to 350 ul of lysed sample in RLT and the mixture was transferred to the RNeasy mini column. RNA was extracted following the manufacturer’s instructions and RNA was eluted from the column with 50 ul of RNase-free water. 2.5 ul of RNA was amplified using the iTaq Universal SYBR Green One Step kit (Biorad) and the following primers (SD29629:AGAAAACGCCGGTAGCAGAA and SD30197:CCTTCCCGAGCCTTCAACAT) using a Biorad CFX Connect Real-time System. The reaction conditions were 10 min at 50C; 5 min at 95C; and 40 cycles of 10 sec at 95C, 20 sec at 59.1C and 36 sec at 72C. All assays contained negative and positive controls. PCR positivity was defined as a rat having a Cq of < 40 for at least 1 observation

### Serology

10 days to six weeks after exposure to SDAV or RCV, animals were bled to assess seroconversion. Animals were restrained in a hand towel and bled from the saphenous vein following shaving, alcohol disinfection of skin, application of sterile petroleum lubricant to create a hydrophobic surface, and single puncture with a 20G needle. Drops of blood were collected in a serum separator tube, spun at 14000 rpm for 10 minutes, and serum withdrawn for storage at −20°C. Sera was tested for coronaviral antibodies using an indirect immunofluorescence assay [36]. Briefly 20 ul of sera diluted tenfold in PBS was placed on a glass slide containing fixed L2p176 cells infected with mouse hepatitis virus strain S. Bound rat antibodies were detected with FITC-conjugated goat anti-rat IgG (Jackson ImmunoResearch).

### Data analysis and statistics

Descriptive statistics were conducted using t-tests, non-parametric tests of medians, and chi-squares tests for proportions where appropriate with data analyzed using regression models. Continuous outcomes (e.g Cq or body weight) used standard linear models with an autoregressive covariance parameter to control for the repeated measurements within animal while count outcomes used Poisson regression with a log-link.

## Results

### Initial infection with SDAV (Table 1)

Body weight, viral shedding and seroconversion were compared across exposure modes.

#### Body weight

Compared to mock-inoculated animals, SDAV-inoculated rats experienced declines in weight gain at days 2-4 post infection (**Supplementary Fig 1**). Weight gain slowed during days 2 – 4 regardless of exposure mode; however, only inoculated rats gained significantly less quickly than mock inoculated control animals (p<.001). Male and female rats experienced no significant difference in growth rates. Apart from transient porphyrin staining of eyes lasting less than 24 hours in 5 rats, no other clinical signs were noted. These data are consistent with clinically asymptomatic to mild infection across all virus exposed groups.

#### Viral shedding

All SDAV-inoculated rats and those exposed via direct contact tested PCR positive on oral swabs (**Table 1**), with declining proportions of PCR-positivity in fomite – exposed animals that were co-housed (73.3%) or singly housed (24%), suggesting that subsequent rat-rat transmission occurred in co-housed animals. Route of exposure significantly influenced amount of viral shedding (p<0.0001; **Supplementary Figure 2**). The extent of viral shedding did not differ between inoculated and direct contact groups. Compared to shedding in inoculated rats, viral shedding following fomite exposure was significantly lower (p< .0001)

**Table 1:** Viral shedding and seroconversion following initial infection with SDAV.

| *Exposure type* | *Duration of exposure* | *PCR positive* | *Cq (mean, range)* | *Shedding time points (mean, range)* | *Sero+* | *Sero-* | *PCR Neg, Sero+* | *PCR Pos, Sero-* |
| --- | --- | --- | --- | --- | --- | --- | --- | --- |
| SDAV 2X10<sup>4</sup> pfu (n=19)* | Inoculation | 19(100%) | 31.5; 25.2-37.4 | 3.1; 2-4 | 19 (100%) | 0 | 0 | 0 |
| Direct contact (n= 31) | 24 hours | 31 (100%) | 31.3; 25.5-36.0 | 2.9; 1-3 | 31 (100%) | 0 | 0 | 0 |
| Fomite-cohab (n=30) | 24 hours | 22 (73.3%) | 32.9; 27.4-36.7 | 1.5; 1-3 | 15 (50%) | 15 (50%) | 9 (30%) | 9 (30%) |
| Fomite-single (n=25) | 24 hours | 6 (24%) | 33.2; 28.6-36.2 | 1.5;1-4 | 2 (8%) | 22 (88%)* | 1 (5%) | 4 (16%) |
| Media inoculation (n=10) | Inoculation | 0 (0%) | Neg, all >40 | 0 | 0 | 10 (100%) | 0 | 0 |

Cq=quantification cycle. Only animals with viral shedding (Cq<40 cycles) are included in Cq and shedding time calculations. Observation time points comprised post-exposure days 2, 3, 4 ,7, 10.
\*One rat in the inoculation group died of unrelated causes before blood was taken for serology

Route of exposure significantly influenced duration of viral shedding (defined as the number of observations, in days, where shedding was present; p<0.0001; **Supplementary Figure 3**). Inoculated and direct exposure groups shed virus for approximately twice as long as fomite exposed animals (significant at p<0.0001 for fomite-cohabitation group only). Inoculated and direct contact animals shed virus from 2-4 days post inoculation, with few animals shedding at day 7 and none at day 10. Fomite exposed animals shed intermittently; however, shedding more commonly persisted to 10 dpi (**Supplementary Figure 4**). Consistency of shedding was significantly affected by exposure mode (p<.0001), with inoculated and direct exposure groups shedding with significantly greater consistency than fomite-cohabitation (p<.0001) and fomite single groups (p<.0001). These data indicate that inoculated and direct contact animals shed higher amounts of virus more consistently in the first 3 days post exposure. In contrast, fomite-exposed animals shed lower amounts of virus at intermittently for a more protracted period.

#### Seroconversion

All SDAV-inoculated rats and those exposed via direct contact seroconverted. Seroconversion declined in cohabiting and singly housed fomite-exposed animals (50% and 8% respectively). These two exposure groups also experienced discordant results between PCR testing and seroconversion (**Table 1**). These data indicate that viral shedding detectable by PCR does not invariably result in seroconversion. Similarly, because viral shedding is intermittent, a negative PCR test does not rule out the potential of infection sufficient to induce an immune response. However, seroconversion was significantly associated with greater amounts (p<0.0001) and duration (p<0.0001) of viral shedding. (**Supplementary Fig 5**). Sex did not significantly affect viral shedding amount, duration or seroconversion.

#### Re-infection with SDAV

Rats were aged for 113-165 days before reinfection with SDAV by inoculation, direct contact or fomite exposure (**Table 2**). In contrast to initial infection, no significant effect on body weight was noted regardless of mode of infection or serologic status. Amount of viral shedding (expressed as lowest observed Cq) was significantly influenced by both exposure mode (p<.01) and serologic status (p<.001; **Figure 3A**). Similarly, duration of shedding was significantly influenced by both exposure mode (p<.05) and serologic status (p<.001; **Figure 3B**).

**Table 2:** Viral shedding after SDAV reinfection in seropositive and seronegative groups

| **A. Seropositive (n=66)** |  |  |  |  |  |  |  |  |  |
| --- | --- | --- | --- | --- | --- | --- | --- | --- | --- |
| *Re-exposure type* | *Duration between infections* | *Duration of exposure* | *PCR pos* | *Cq (mean, range)* | *Shedding time points (mean, range)* | *Sero+* | *Sero-* | *PCR Neg, Sero+* | *PCR Pos, Sero-* |
| SDAV 2X10<sup>4</sup> pfu intranasal (n=18)* | 113-144 days | Inoculation | 7(38.9%) | 32.7; 28.9-36.4 | 1.7;1-4 | - | - | - | - |
| Direct contact (n=19) | 143-165 days | 24 hours | 15 (78.9%) | 34.1; 29.1-36.6 | 2; 1-4 | - | - | - | - |
| Fomite-cohab (n=17) | 114-143 days | 24 hours | 13 (76.5%) | 34.4; 30.7-37.9 | 1.7; 1-2 | - | - | - | - |
| Fomite-single (n=12) | 114-165 days | 24 hours | 4 (33.3%) | 34.5; 33.0-36.2 | 1.7; 1 | - | - | - | - |
| **A. Seronegative (n=40)** |  |  |  |  |  |  |  |  |  |
| SDAV 2X10<sup>4</sup> pfu intranasal (n=18)** | 82-163 days | Inoculation | 18 (100%) | 30.6; 23.5-35.3 | 3; 2-4 | 18 (100%) | 0 | 0 | 0 |
| Direct contact (n=9) | 143-165 days | 24 hours | 8 (88.8%) | 31.9; 27.3-33.4 | 1.4; 1-3 | 8 (88.8%) | 1 (11.1%) | 1 (11.1%) | 1 (11.1%) |
| Fomite-cohab (n=9) | 165 days | 24 hours | 6 (66.6%) | 31.3; 27.2-34.5 | 1.6;1-4 | 9 (100%) | 0 | 0 | 0 |
| Fomite-single (n=4) | 165 days | 24 hours | 1 (25%) | 33.9; 33.9-34.0 | 2; 2 | 1 (25%) | 3 (75%) | 1 (25%) | 3 (75%) |

\*Seropositive group: Animals given SDAV 2X10<sup>4</sup> pfu on initial infection were reinfected via the same route. All other animals were randomized between initial and subsequent routes of infection and were exposed to naïve animals given SDAV 2X10<sup>4</sup> pfu via direct contact or fomite exposure.
\*\*Seronegative group: Animals given SDAV 2X10<sup>4</sup> pfu reinfection were previously mock infected controls or naïve purchased animals. All other animals were exposed to naïve animals inoculated with SDAV 2X10<sup>4</sup> pfu via direct contact or fomite exposure.
- Seropositive animals were not re-tested for seroconversion after reinfection
Cq=quantification cycle. Only animals with viral shedding (Cq<40 cycles) are included in Cq and shedding time calculations. Observation time points comprised post-exposure days 2, 3, 4, 7, 10.

**Table 3:** Transmission of SDAV by previously SDAV infected or RCV vaccinated animals

| **A. Transmission after SDAV reinfection** |  |  |  |  |  |  |  |  |  |  |
| --- | --- | --- | --- | --- | --- | --- | --- | --- | --- | --- |
| Source animals (n=13) |  |  |  |  | Susceptible recipient animals (n=13) |  |  |  |  |  |
| *Initial infection* | *Re-infection* | *Duration between infections* | *PCR pos* | *Cq; shed time points (mean; range)* | *Duration of exposure* | *PCR pos* | *Cq; time points shedding* | *Sero+* | *PCR Neg, Sero+* | *PCR Pos, Sero-* |
| SDAV 2X10<sup>4</sup> pfu | SDAV 2X10<sup>4</sup> pfu | 113-140 days | 2 (15.4%) | 32.2; 29.2-34.4<br>1.5; 1-2 | 24 hours | 3 (23.1%) | 34.0; 32.7-36.5<br>2.3; 1-4 | 3 (23.1%) | 1 (7.7%) | 1 (7.7%) |
| **B. Transmission after RCV single vaccination** |  |  |  |  |  |  |  |  |  |  |
| Source animals (n=12) |  |  |  |  | Susceptible recipient animals (n=12) |  |  |  |  |  |
| RCV 10<sup>3</sup> pfu | SDAV 2X10<sup>4</sup> pfu | 48 days | 4 (33.3%) | 32.5; 30.1-34.5<br>1.8; 1-3 | 7 days | 1 (8.3%) | 31.8; 28.5-25.5<br>3; 3 | 1 (8.3%) | 0 | 0 |
| **C. Transmission after RCV double vaccination** |  |  |  |  |  |  |  |  |  |  |
| Source animals (n=12) |  |  |  |  | Susceptible recipient animals (n=9) |  |  |  |  |  |
| RCV 10e3 pfu twice, 30 days apart | SDAV 2X10<sup>4</sup> pfu | 42 days | 7 (58.3%) | 34.7; 29.5-37.6<br>1.6; 1-3 | 7 days | 2 (22.2%) | 32.5; 32.3-32.9<br>1.5; 1-2 | 0 (0%) | 0 | 0 |

Viral shedding and seroconversion in naïve (susceptible recipient) animals exposed to SDAV inoculated animals.
Cq=quantification cycle. Only animals with viral shedding (Cq<40 cycles) are included in Cq and shedding time calculations. Observation time points comprised post-exposure days 2, 3, 4, 7, 10.

**Figure 3.**
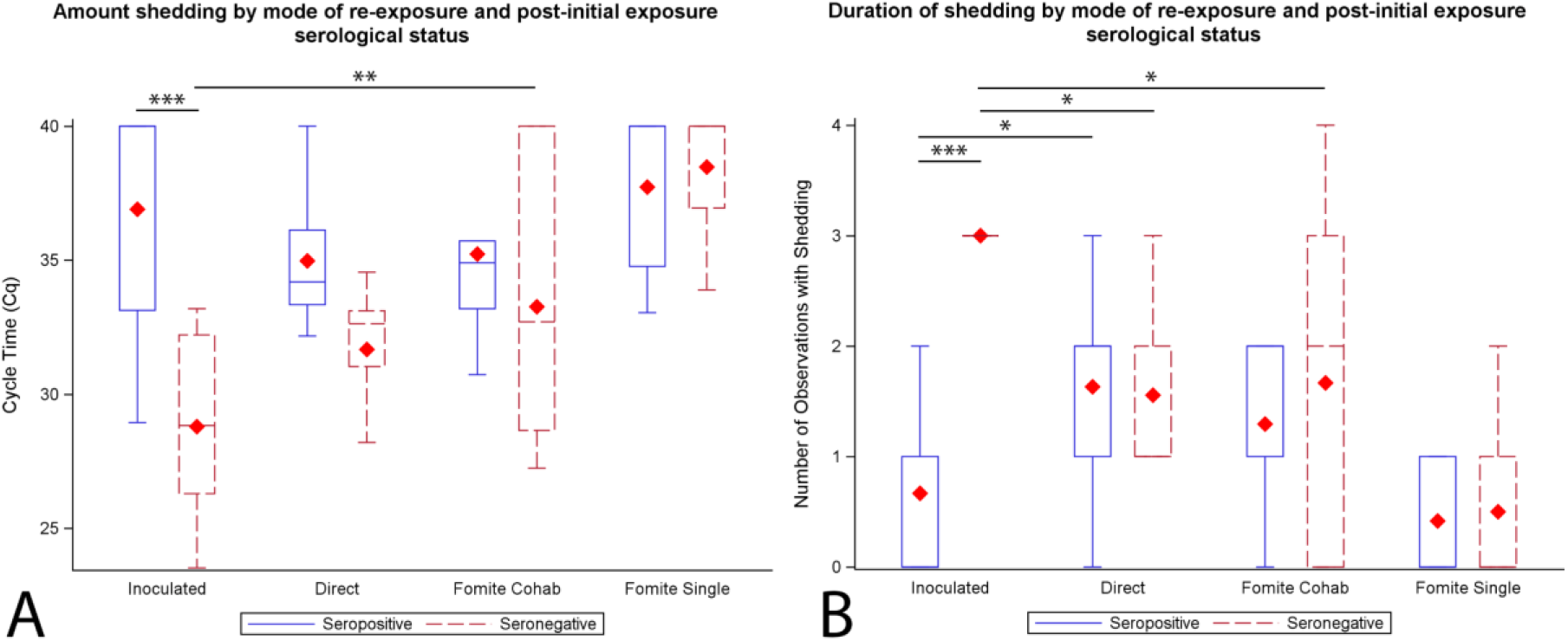
Viral shedding on SDAV re-exposure. A. Amount of viral shedding (expressed as lowest observed Cq) by serologic status and exposure mode. Amount of viral shedding was significantly influenced by both exposure mode (p< .01) and serologic status (p<.001) using linear regression. Previously SDAV-exposed seropositive rats shed less virus (p <.001) compared to previously mock-inoculated seronegative rats. Among seronegative rats, exposure mode significantly affected amount (p < .001) of viral shedding. Compared to seronegative SDAV-inoculated rats, direct contact (p=.06) and fomite cohabitation animals(p<.01) shed less virus. Among seropositive rats, direct and fomite cohabitating rats experienced higher levels of shedding than the SDAV-inoculated seropositive rats, implying greater protection against shedding in the latter. Despite this observation, the amount of viral shedding was not significantly associated with exposure mode (p=.09). Red diamonds indicate group means. Contrasts are marked by asterisks; * p<.05, ** p<.01, *** p<.001. Boxplot of lowest observed Cq over the entire 10-day observation period by exposure mode B. Duration of viral shedding (observed shedding events) by serologic status and exposure mode. Duration of shedding was significantly influenced by both exposure mode (p< .05) and serologic status (p<.001) using Poisson regression. Previously SDAV-exposed seropositive rats shed virus for a shorter period (p <.001) compared to previously mock-inoculated seronegative rats when re-inoculated with the same dose of SDAV. Among seronegative rats, exposure mode significantly affected duration (p < .01) of viral shedding. Compared to seronegative SDAV-inoculated rats, direct contact rats shed virus for a fewer observations (p<0.05), as did fomite cohabitation animals (p<0.05). Among seropositive rats, overall exposure mode was significantly (p<.01) associated with duration of shedding. SDAV-inoculated rats generally shed for fewer observations, and significantly fewer observations the direct contact group (p<.05). Red diamonds indicate group means. Contrasts are marked by asterisks; * p<.05, ** p<.01, *** p<.001.

Previously SDAV-exposed seropositive rats shed less virus (p<.001) for a shorter period (p <.001) compared to previously mock-inoculated seronegative rats when re-inoculated with the same dose of SDAV. Proportions infected among SDAV inoculated rats differed significantly (P<.0001) between seronegative (100%) and seropositive rats (38.9%). This clearly demonstrates protective immunity (that did not however eliminate viral shedding) on reinfection with the same virus and same dose.

To investigate the differential role of exposure mode on amount of viral shedding, rats were stratified by serological status (**Table 2**). Among seronegative rats, exposure mode significantly affected amount (p<.001) and duration (p < .0001) of viral shedding. Compared to seronegative SDAV-inoculated rats, direct contact rats shed less virus (p=.06) for a shorter period of time (p<0.001) Fomite cohabitation animals shed less virus (p<.01) for a shorter period of time (p<0.001). The fomite single group was excluded from analysis due to low sample size (n=4). Among seronegative animals, as with initial infection, the amount and duration of viral shedding was dictated by exposure mode; however, direct contact rats appeared to shed less virus and for less time than on initial infection.

In seropositive animals, immunity obtained from prior SDAV exposure via a variety of routes altered this pattern. Direct and fomite cohabitating rats experienced higher levels of shedding than the SDAV-inoculated seropositive rats previously exposed to SDAV via the same route, implying greater protection against shedding in the latter. Despite this observation, the amount of viral shedding was not significantly associated with exposure mode (p=.09). In contrast, exposure mode was significantly (p<.01) associated with duration of shedding. SDAV-inoculated rats generally shed for fewer observations, and significantly fewer observations than the direct contact group (p<.05). These data indicate that immunity obtained via direct contact or fomite exposure was largely protective regardless of how it was obtained, but that heterogeneity in protection against duration of shedding was imparted by route of re-exposure.

#### Viral transmission by SDAV reinfected or RCV vaccinated rats (**Table 3**)

In the previous experiment, we established that the majority (59%) of seropositive animals could shed virus, albeit at low levels, on re-exposure. Next, we explored whether such animals could transmit infection to naïve animals. First, we assessed whether rats reinfected with SDAV via the same route (intranasal inoculation of 2X10^e^4 pfu/animal) could transmit the virus to naïve animals using the same 24-hour direct contact exposure paradigm employed in our previous experiments. The time-period between initial SDAV infection and reinfection was 113-140 days, thus modeling shedding on re-exposure after several months. The same experiment was repeated following one or two doses of RCV given one month apart. In this experiment, direct contact exposure times were lengthened to 7 days to determine if SDAV transmission could occur at all. Following SDAV inoculation, no significant differences in number of animals shedding virus, amount of virus shed or duration of shedding were noted across vaccine groups, indicating that one or two doses provided equivalent protection. Transmission was defined in two ways 1) if the target rats exhibited shedding on any observation and 2) if the target rat seroconverted. Transmission rates from the SDAV reinfected (n=13) and RCV vaccinated rats (n=21, collapsing 1 and 2 doses) were compared to each other and to a reference group consisting of the previously described rats in direct contact with SDAV naïve rats (n=31) using chi-square tests. Regardless if shedding or seroconversion was treated as the metric of transmission, proportions in the both SDAV reinfected source group (3/13, shedding and seroconversion) and RCV vaccinated group (3/21, shedding; 1/21, seroconversion) were significantly lower than the SDAV naïve direct contact reference group (31/31, shedding and seroconversion), p<.001 in both cases. However, the proportions did not differ between SDAV reinfected and RCV vaccinated SDAV exposed rats (shedding p=.65; seroconversion, p=.27).

These data indicate that low levels of viral shedding can occur in previously infected and vaccinated animals on SDAV re-exposure and challenge respectively. These levels, while low, can result in transmission, even in the period following vaccination, when immunity is likely to be the highest. However, rates of transmission by previously infected and vaccinated animals are far lower than those seen with initial infection.

## Discussion

The extent to which humans with natural or vaccine-induced immunity who are re-exposed to SARS-CoV-2 can shed virus is only just emerging [12]. Immunity following natural infection is likely to be variable [13]. Vaccine-induced immune protection is more homogenous [37,38], however efficacy varies across different vaccines [8] and is likely to be impacted by newly emerging variants [39,40]. This immunologic heterogeneity may promote persistent viral transmission and continued disease in remaining susceptible individuals. A key issue is whether low amounts of shed virus can achieve transmission [41]. Using a rat model of respiratory coronaviral infection, we modeled viral shedding and transmission in a population exposed via routes varying from high (inoculation and direct contact) to low (fomite) exposure risk. Of particular interest was the extent to which heterogeneous immune protection afforded by prior exposure influenced the amount of virus shed on re-exposure, and whether these low amounts of shed virus could infect naïve animals.

SDAV is a highly infectious rat betacoronavirus [17,18] that infects the upper respiratory tract [19], lacrimal glands [20,21] and lung [21–23], and results in transient mild to asymptomatic disease. SDAV infects salivary glands [21], a capacity recently demonstrated for SARS-CoV-2 [42]. Both SARS-CoV-2 and SDAV can persist on hard surfaces for up to 28 days [27,43] and 2 days [44] respectively. Like SARS CoV-2 [24–28], SDAV infection can be transmitted by asymptomatically infected individuals [29] via airborne, direct contact or fomite routes [17].

Consequently, SDAV is well suited to model viral transmission dynamics of COVID-19 in asymptomatic people [24], the predominant means of community spread.

On initial infection, animals exposed via all routes experienced transient declines in growth rate, consistent with mild to asymptomatic disease. This was significant only in inoculated rats, who received a known (and presumably highest) infective dose. Like SARS-CoV-2, viral shedding peaked early in the first week after exposure [45], however duration of viral shedding (3-10 days) in SDAV infection was shorter than the reported mean of 17.5 days in the upper respiratory tract of SARS-CoV2 infected patients [45]. Amount and duration of viral shedding was significantly influenced by route of exposure, and was highest in inoculated and direct contact groups, and lowest fomite groups. Fomite transmission was amplified by subsequent cohabitation of rats. Conflicting data exist regarding the risk of fomite transmission in SARS-CoV-2 [27,46]. Our data indicate that while comparatively low, this route represents a clear source of transmission risk in that can be amplified by close contact such as dense co-housing conditions.

As in the human population, the viral dose that resulted in infection of an individual cannot be directly measured in natural exposure settings. However, higher infectious doses of SARS-CoV-2 are implicated in higher risk of transmission [47], and infectious dose is assumed to be much higher in direct contact compared to fomite settings, corresponding to respective high and low transmission risk of these exposure types [48,49]. Similarly, higher viral loads in COVID-19 patients are associated with more reliable seroconversion [50]. Our results are consistent with these data and imply that higher viral exposure results in a greater amount and duration of viral shedding, and more consistent seroconversion. Conversely, low viral load accompanying with fomite exposure may result in a positive viral PCR test but fail to elicit an antibody response.

Following initial infection with SDAV, we generated a population of seronegative and seropositive animals. Within this latter group, 59% of animals shed virus for some period on re-exposure, although at lower levels than in initial infection. Those animals with the highest confirmed viral exposure on initial infection and re-infection (i.e. intranasal inoculation for both exposures) had significantly lower viral shedding for a shorter period than all other reinfection groups. The remaining seropositive rats achieved their status through initial exposure via direct contact or fomite exposure where viral dose was unknown, but presumably lower than that used for direct inoculation. On reinfection via one of these routes, re-exposed rats shed virus for a longer period of time, indicating that their protective immunity was lower than that induced by intranasal inoculation. These results indicate that seropositive status following initial infection reduces but does not eliminate viral shedding on re-exposure after 3.7-5.5 months. Moreover, immune protection in seropositive individuals appears to be heterogeneous. The extent to which humans shed SARS-CoV-2 on reinfection, and associated risk of transmission is emerging [51]. Shedding rates on reinfection in our model are much higher than those reported for SARS-CoV-2 [9]. This may reflect a true biological difference between SDAV and SARS-CoV-2. However, our entire population was re-exposed within a defined time-period and shedding then assessed by timed repeated PCR testing, thus maximizing the likelihood of detecting shedding. Re-exposure events in a human population are rarely known, thus precluding coordination of testing with the exposure event. Therefore, the actual shedding on re-exposure in human populations may be higher than reported.

Next, we assessed the risk of transmission by animals shedding low amounts of virus. Consistent with prior studies [52,53], rats reinfected with SDAV via the same route (intranasal inoculation given 113-140 days apart) were able to induce viral shedding and/or seroconversion in 25% of naïve contact rats after 24-hour exposure by direct contact. Observed cycle threshold values in reinfected source rats ranged from 29-34 cycles with short shedding durations (1-2 days). In SARS-CoV-2, live viral shedding can be inferred from cycle threshold (Cq) values on PCR testing [45] of shedding individuals. Live viral culture positivity declines with increasing cycle threshold values [54]. Cycle threshold values of 33-34 reflect low enough live viral shedding to render patients non-contagious [25,54,55]. These reports are consistent with our findings. However, it should be noted that in all three instances where naïve recipient rats seroconverted, viral shedding by SDAV re-infected rats was not detected, implying that shedding sufficient for transmission can occur for short periods that may escape detection. While both SARS-CoV-2 [42] and SDAV infect salivary glands and can be detected in the oral cavity, we recognize that oral swabs in our rats may not have detected shedding via other routes e.g. nasal shedding. Implications for COVID-19 are that naturally infected seropositive individuals may represent a source of ongoing transmission on re-exposure to SARS-CoV-2.

Finally, we assessed transmission by animals exposed to SDAV shortly after vaccination with Parker’s Rat Coronavirus (RCV), a betacoronaviruses that is closely related to SDAV [31]. RCV infection can be used to model vaccination in our model using a live virus strategy. In prior studies, infection with RCV results in cross-protective seroconversion, and disease protection following subsequent SDAV infection [56]. While a significant proportion of vaccinated animals (11/24) shed SDAV at low amounts (range 29.5-36 cycles), they achieved transmission in only one recipient after 7 days of direct contact exposure. These results indicate that protection against transmission after reinfection in the immediate post vaccination period is superior to that several months after natural infection. We did not test whether this protection would decline over months at the same rate as that afforded by SDAV inoculation, however some decline is expected from previous studies [56]. Data from both groups taken together imply that cycle threshold (Cq) values above 29 are associated with transmission, but that PCR detection of oral shedding alone is not a reliable predictor of transmission risk.

In conclusion, shedding and seroconversion following initial natural SDAV infection was heterogeneous and influenced by route of exposure. Viral shedding occurred in the majority seropositive individuals exposed to SDAV after natural infection or vaccination, however amount and duration of shedding was much lower than on initial exposure. Nevertheless, low viral shedding by these animals was able to result in transmission to susceptible individuals after close contact. If these data are extrapolated to SARS-CoV-2 transmission, it appears that viral shedding and transmission by previously infected or vaccinated individuals could prolong transition to stable endemic status. This transition could be impacted by emergence of more transmissible variants. Protection of susceptible individuals would best be achieved by vaccination rather than relying on protection by herd immunity.

## Acknowledgements

The authors thank members of the Yale Animal Resources Center for their excellent animal care.

## References

1. Alwan NA, Burgess RA, Ashworth S, et al. Scientific consensus on the COVID-19 pandemic: we need to act now. Lancet. 2020 Oct 31;396(10260):e71–e72.

2. King C, Einhorn L, Brusselaers N, et al. COVID-19-a very visible pandemic. Lancet. 2020 Aug 8;396(10248):e15.

3. Omer SB, Yildirim I, Forman HP. Herd Immunity and Implications for SARS-CoV-2 Control. JAMA: the journal of the American Medical Association. 2020 Nov 24;324(20):2095–2096.

4. Lai CC, Wang JH, Hsueh PR. Population-based seroprevalence surveys of anti-SARS-CoV-2 antibody: An up-to-date review. Int J Infect Dis. 2020 Dec;101:314–322.

5. Manisty C, Treibel TA, Jensen M, et al. Time series analysis and mechanistic modelling of heterogeneity and sero-reversion in antibody responses to mild SARS-CoV-2 infection. EBioMedicine. 2021 Mar;65:103259.

6. Long COVID: let patients help define long-lasting COVID symptoms. Nature. 2020 Oct;586(7828):170.

7. Woolf SH, Chapman DA, Lee JH. COVID-19 as the Leading Cause of Death in the United States. JAMA: the journal of the American Medical Association. 2021 Jan 12;325(2):123–124.

8. Anderson RM, Vegvari C, Truscott J, et al. Challenges in creating herd immunity to SARS-CoV-2 infection by mass vaccination. Lancet. 2020 Nov 21;396(10263):1614–1616.

9. Hansen CH, Michlmayr D, Gubbels SM, et al. Assessment of protection against reinfection with SARS-CoV-2 among 4 million PCR-tested individuals in Denmark in 2020: a population-level observational study. Lancet. 2021 Mar 27;397(10280):1204–1212.

10. Adrielle Dos Santos L, Filho PGG, Silva AMF, et al. Recurrent COVID-19 including evidence of reinfection and enhanced severity in thirty Brazilian healthcare workers. J Infect. 2021 Mar;82(3):399–406.

11. Yang Y, Wang X, Du RH, et al. Serological investigation of asymptomatic cases of SARS-CoV-2 infection reveals weak and declining antibody responses. Emerg Microbes Infect. 2021 Apr 19:1–28.

12. Levine-Tiefenbrun M, Yelin I, Katz R, et al. Initial report of decreased SARS-CoV-2 viral load after inoculation with the BNT162b2 vaccine. Nature medicine. 2021 Mar 29.

13. Britton T, Ball F, Trapman P. A mathematical model reveals the influence of population heterogeneity on herd immunity to SARS-CoV-2. Science (New York, NY). 2020 Aug 14;369(6505):846–849.

14. Wajnberg A, Amanat F, Firpo A, et al. Robust neutralizing antibodies to SARS-CoV-2 infection persist for months. Science (New York, NY). 2020 Dec 4;370(6521):1227–1230.

15. Iyer AS, Jones FK, Nodoushani A, et al. Persistence and decay of human antibody responses to the receptor binding domain of SARS-CoV-2 spike protein in COVID-19 patients. Sci Immunol. 2020 Oct 8;5(52).

16. Marot S, Malet I, Leducq V, et al. Rapid decline of neutralizing antibodies against SARS-CoV-2 among infected healthcare workers. Nature communications. 2021 Feb 8;12(1):844.

17. La Regina M, Woods L, Klender P, et al. Transmission of sialodacryoadenitis virus (SDAV) from infected rats to rats and mice through handling, close contact, and soiled bedding. Laboratory animal science. 1992 Aug;42(4):344–6.

18. Bhatt PN, Percy DH, Jonas AM. Characterization of the virus of sialodacryoadenitis of rats: a member of the coronavirus group. The Journal of infectious diseases. 1972 Aug;126(2):123–30.

19. Bihun CG, Percy DH. Morphologic changes in the nasal cavity associated with sialodacryoadenitis virus infection in the Wistar rat. Veterinary pathology. 1995 Jan;32(1):1–10.

20. Eisenbrandt DL, Hubbard GB, Schmidt RE. A subclinical epizootic of sialodacryoadenitis in rats. Laboratory animal science. 1982 Dec;32(6):655–9.

21. Percy DH, Hanna PE, Paturzo F, et al. Comparison of strain susceptibility to experimental sialodacryoadenitis in rats. Laboratory animal science. 1984 Jun;34(3):255–60.

22. Wojcinski ZW, Percy DH. Sialodacryoadenitis virus-associated lesions in the lower respiratory tract of rats. Veterinary pathology. 1986 May;23(3):278–86.

23. Funk CJ, Manzer R, Miura TA, et al. Rat respiratory coronavirus infection: replication in airway and alveolar epithelial cells and the innate immune response. The Journal of general virology. 2009 Dec;90(Pt 12):2956–2964.

24. Johansson MA, Quandelacy TM, Kada S, et al. SARS-CoV-2 Transmission From People Without COVID-19 Symptoms. JAMA Netw Open. 2021 Jan 4;4(1):e2035057.

25. Arons MM, Hatfield KM, Reddy SC, et al. Presymptomatic SARS-CoV-2 Infections and Transmission in a Skilled Nursing Facility. The New England journal of medicine. 2020 Apr 24.

26. Zhang R, Li Y, Zhang AL, et al. Identifying airborne transmission as the dominant route for the spread of COVID-19. Proceedings of the National Academy of Sciences of the United States of America. 2020 Jun 30;117(26):14857–14863.

27. Kraay ANM, Hayashi MAL, Berendes DM, et al. Risk for Fomite-Mediated Transmission of SARS-CoV-2 in Child Daycares, Schools, Nursing Homes, and Offices. Emerg Infect Dis. 2021 Apr;27(4):1229–1231.

28. Hu S, Wang W, Wang Y, et al. Infectivity, susceptibility, and risk factors associated with SARS-CoV-2 transmission under intensive contact tracing in Hunan, China. Nature communications. 2021 Mar 9;12(1):1533.

29. Bhatt PN, Jacoby RO. Epizootiological observations of natural and experimental infection with sialodacryoadenitis virus in rats. Laboratory animal science. 1985 Apr;35(2):129–34.

30. Parker JC, Cross SS, Rowe WP. Rat coronavirus (RCV): a prevalent, naturally occurring pneumotropic virus of rats. Arch Gesamte Virusforsch. 1970;31(3):293–302.

31. Barker MG, Percy DH, Hovland DJ, et al. Preliminary characterization of the structural proteins of the coronaviruses, sialodacryoadenitis virus and Parker’s rat coronavirus. Can J Vet Res. 1994 Apr;58(2):99–103.

32. Procop GW, Shrestha NK, Vogel S, et al. A Direct Comparison of Enhanced Saliva to Nasopharyngeal Swab for the Detection of SARS-CoV-2 in Symptomatic Patients. Journal of clinical microbiology. 2020 Oct 21;58(11).

33. Mohit E, Rostami Z, Vahidi H. A comparative review of immunoassays for COVID-19 detection. Expert Rev Clin Immunol. 2021 Mar 31.

34. Gaertner DJ, Winograd DF, Compton SR, et al. Development and optimization of plaque assays for rat coronaviruses. J Virol Methods. 1993 Jun;43(1):53–64.

35. Gaertner DJ, Smith AL, Paturzo FX, et al. Susceptibility of rodent cell lines to rat coronaviruses and differential enhancement by trypsin or DEAE-dextran. Arch Virol. 1991;118(1-2):57–66.

36. Smith AL. An immunofluorescence test for detection of serum antibody to rodent coronaviruses. Laboratory animal science. 1983 Apr;33(2):157–60.

37. Walsh EE, Frenck RW, Jr., Falsey AR, et al. Safety and Immunogenicity of Two RNA-Based Covid-19 Vaccine Candidates. The New England journal of medicine. 2020 Dec 17;383(25):2439–2450.

38. Folegatti PM, Ewer KJ, Aley PK, et al. Safety and immunogenicity of the ChAdOx1 nCoV-19 vaccine against SARS-CoV-2: a preliminary report of a phase 1/2, single-blind, randomised controlled trial. Lancet. 2020 Aug 15;396(10249):467–478.

39. Abdool Karim SS, de Oliveira T. New SARS-CoV-2 Variants - Clinical, Public Health, and Vaccine Implications. The New England journal of medicine. 2021 Mar 24.

40. Altmann DM, Boyton RJ, Beale R. Immunity to SARS-CoV-2 variants of concern. Science (New York, NY). 2021 Mar 12;371(6534):1103–1104.

41. Widders A, Broom A, Broom J. SARS-CoV-2: The viral shedding vs infectivity dilemma. Infect Dis Health. 2020 Aug;25(3):210–215.

42. Huang N, Pérez P, Kato T, et al. SARS-CoV-2 infection of the oral cavity and saliva. Nature medicine. 2021 Mar 25.

43. van Doremalen N, Bushmaker T, Morris DH, et al. Aerosol and Surface Stability of SARS-CoV-2 as Compared with SARS-CoV-1. The New England journal of medicine. 2020 Apr 16;382(16):1564–1567.

44. Gaertner DJ, Compton SR, Winograd DF. Environmental stability of rat coronaviruses (RCVs). Laboratory animal science. 1993 Oct;43(5):403–4.

45. Cevik M, Tate M, Lloyd O, et al. SARS-CoV-2, SARS-CoV, and MERS-CoV viral load dynamics, duration of viral shedding, and infectiousness: a systematic review and meta-analysis. Lancet Microbe. 2021 Jan;2(1):e13–e22.

46. Lewis D. COVID-19 rarely spreads through surfaces. So why are we still deep cleaning? Nature. 2021 Feb;590(7844):26–28.

47. Kawasuji H, Takegoshi Y, Kaneda M, et al. Transmissibility of COVID-19 depends on the viral load around onset in adult and symptomatic patients. PloS one. 2020;15(12):e0243597.

48. Wodarz D, Komarova NL, Schang LM. Role of high-dose exposure in transmission hot zones as a driver of SARS-CoV-2 dynamics. J R Soc Interface. 2021 Mar;18(176):20200916.

49. Jones RM. Relative contributions of transmission routes for COVID-19 among healthcare personnel providing patient care. J Occup Environ Hyg. 2020 Sep;17(9):408–415.

50. Masiá M, Telenti G, Fernández M, et al. SARS-CoV-2 Seroconversion and Viral Clearance in Patients Hospitalized With COVID-19: Viral Load Predicts Antibody Response. Open Forum Infect Dis. 2021 Feb;8(2):ofab005.

51. Ledford H. Coronavirus reinfections: three questions scientists are asking. Nature. 2020 Sep;585(7824):168–169.

52. Weir EC, Jacoby RO, Paturzo FX, et al. Infection of SDAV-immune rats with SDAV and rat coronavirus. Laboratory animal science. 1990 Jul;40(4):363–6.

53. Percy DH, Bond SJ, Paturzo FX, et al. Duration of protection from reinfection following exposure to sialodacryoadenitis virus in Wistar rats. Laboratory animal science. 1990 Mar;40(2):144–9.

54. La Scola B, Le Bideau M, Andreani J, et al. Viral RNA load as determined by cell culture as a management tool for discharge of SARS-CoV-2 patients from infectious disease wards. Eur J Clin Microbiol Infect Dis. 2020 Jun;39(6):1059–1061.

55. Tom MR, Mina MJ. To Interpret the SARS-CoV-2 Test, Consider the Cycle Threshold Value. Clin Infect Dis. 2020 Nov 19;71(16):2252–2254.

56. Bihun CG, Percy DH. Coronavirus infections in the laboratory rat: degree of cross protection following immunization with a heterologous strain. Can J Vet Res. 1994 Jul;58(3):224–9.

