## Supplemental Figures for "Viral shedding and transmission after natural infection and vaccination in an animal model of SARS-CoV-2 propagation"

Supplementary Figure 1

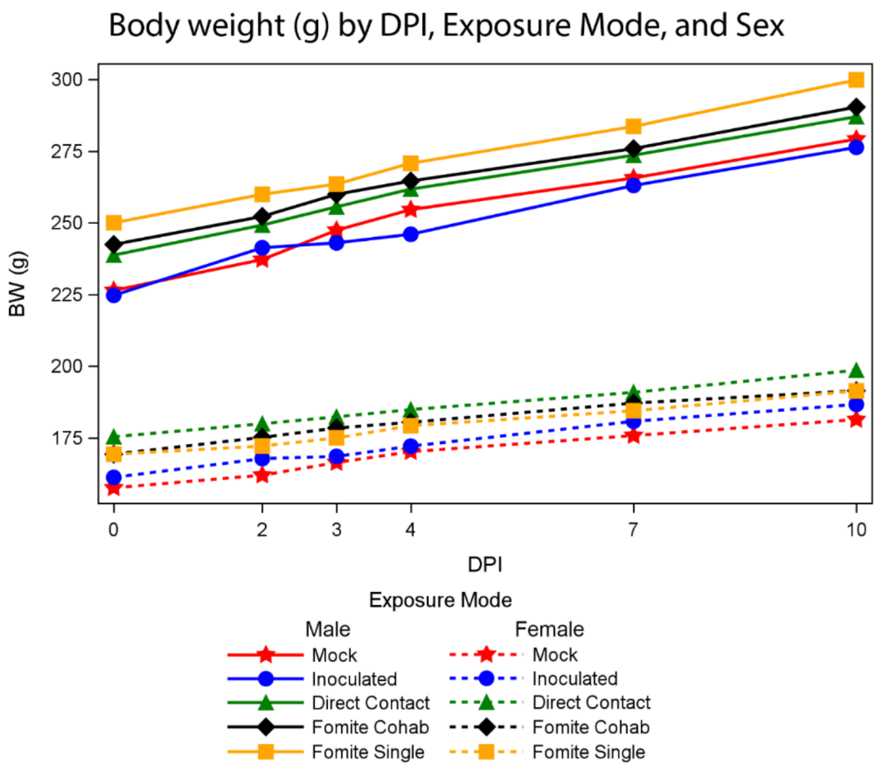


**Supplementary Fig 1**. Body weight by exposure mode, DPI 0-10.

Compared to mock-inoculated animals, SDAV-inoculated rats experienced declines in weight gain at days 2-4 post infection. Weight gain slowed during days 2 – 4 regardless of exposure mode, however, only index rats grew significantly less quickly than mock inoculated control animals ($p<.001$). Male and female rats experienced no significant difference in growth rates.

Body weight was modeled with fixed effects of sex, exposure mode, time, and a mode by time interaction using mixed repeated measures linear models with an autoregressive covariance structure (AR(1)) accounting for correlations of repeated measurement within animal. A second model on the subset of index exposed rats was run to test whether the pronounced slowing of growth differed by sex. But the interaction was not significant, indicating similar slowed growth in male and female rats. Solid lines depict male rats and dashed lines depict female rats.

Supplementary Figure 2

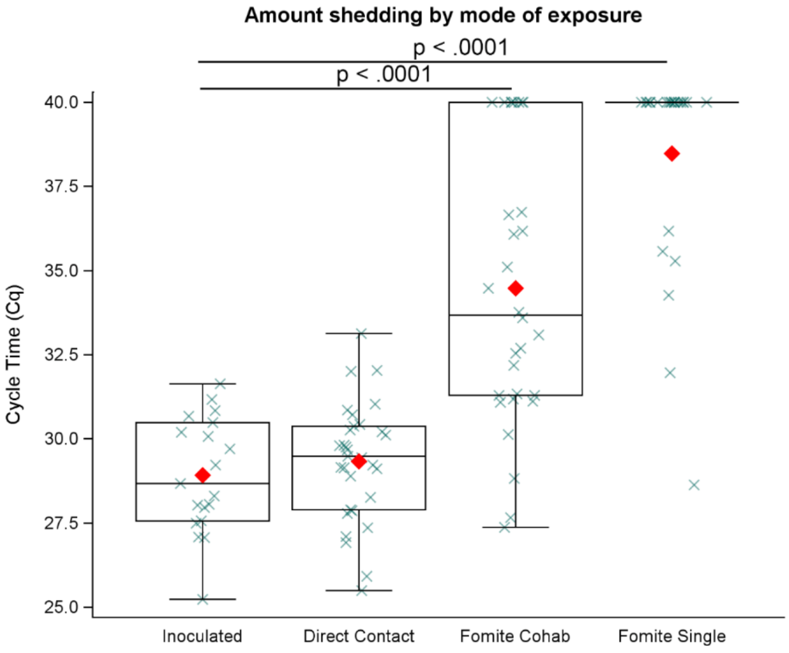


**Supplementary Figure 2.** Amount of viral shedding following initial SDAV exposure.

Route of exposure significantly influenced amount of viral shedding ($p<0.0001$). Extent of viral shedding did not differ significantly between index and direct contact groups. Using the inoculated group as a reference, viral shedding following fomite exposure (both cohabitation and single groups) was significantly lower ($p<.0001$). Linear regression modeled the lowest observed PCR as function of exposure mode, with planned contrasts between exposure modes tested with t-tests. Red diamonds indicate group means. Individual rat data are depicted with green x-marks.

Supplementary Figure 3

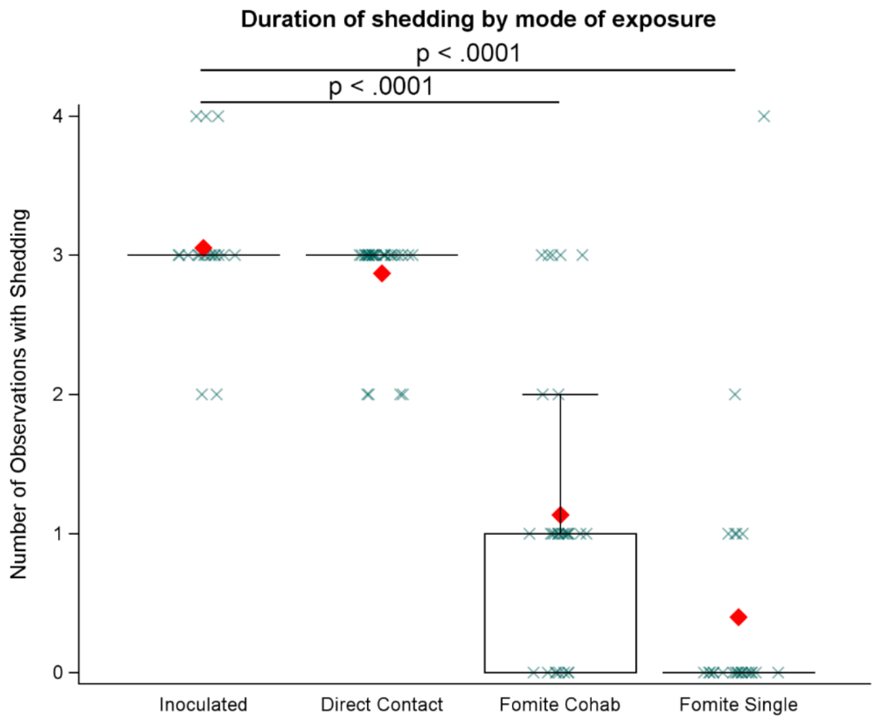


**Supplementary Figure 3.** Duration of viral shedding following initial SDAV exposure.

The count of observations with shedding was modeled with a Poisson linear regression with a log link as a function of exposure mode. Overall, exposure mode was significant ($p<.0001$). Inoculated and direct exposure groups shed virus for approximately twice as long (average 3.1 and 2.9 days respectively) as fomite exposed animals (average 1.5 days, significant at $p<0.0001$ for fomite-cohabitation group compared to inoculation group). Red diamonds indicate group means. Individual rat data are depicted with green x-marks.

Supplementary Figure 4

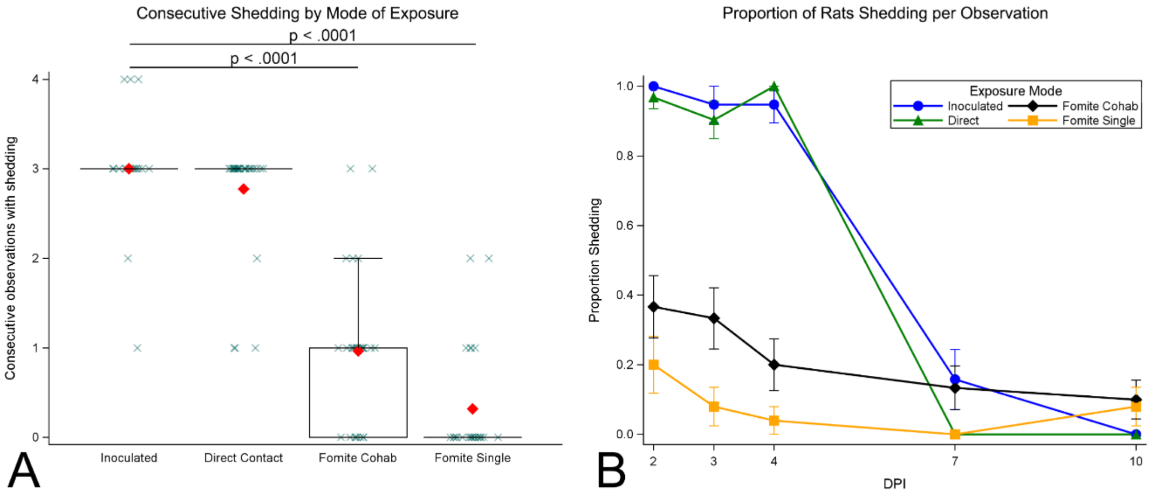


**Supplementary Figure 4:** Proportion of rats shedding per observation time by exposure mode

Inoculated and direct contact animals shed virus from days 2-4 days post inoculation, with few animals shedding at day 7 and none at day 10. Fomite exposed animals shed intermittently, however shedding more commonly persisted to 10 dpi.

**Panel A**: The count of consecutive observations with shedding was modeled with a Poisson linear regression with a log link as a function of exposure mode. Consistency of shedding was significantly affected by exposure mode ($p < .0001$), with inoculated and direct exposure groups shedding with significantly greater consistency than fomite-cohabitation ($p<.0001$) and fomite single groups ($p<.0001$). Red diamonds indicate group means. Individual rat data are depicted with green x-marks.

**Panel B** depicts the proportion of rats shedding within each exposure mode for each observation period.

Supplementary Figure 5

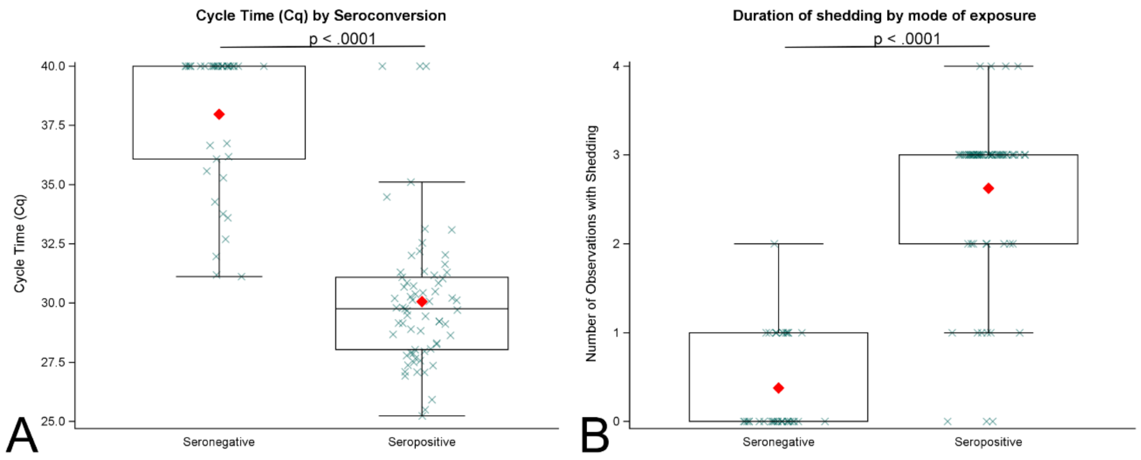


**Supplementary Fig 5:** Amount of viral shedding and seroconversion.

Seroconversion was significantly associated with greater amounts (A; p<.0001) and duration (B; p<.0001) of viral shedding. Sex did not significantly affect viral shedding amount, duration or seroconversion. Differences between shedding as a function of seroconversion was assessed using a t-test on the observed lowest cycle times (Cq). Differences between number of observations with shedding as a function of seroconversion was assessed using a non-parametric test of medians. Red diamonds indicate group means. Individual rat data are depicted with green x-marks.

Boxplot of lowest observed Cq by seroconversion status
